# Employment of self-organisation to achieve economy of scale in biology

**DOI:** 10.1101/2025.03.21.644553

**Authors:** Eric Siero, Eva Slegers, Koen Herman, Robert-Jan Bleichrodt

**Affiliations:** Department of Plant Sciences, Biometris, Wageningen University and Research, Wageningen, The Netherlands; Department of Earth Sciences, Geosciences, Utrecht University, Utrecht, The Netherlands; Microbiology, Department of Biology, Utrecht University, Utrecht, The Netherlands

**Keywords:** Cellular heterogeneity, *Aspergillus niger*, enzymatic degradation, specialisation and optimisation, multi-step processes, compartmentalisation

## Abstract

Self-organisation is a striking phenomenon that fascinates many. Especially in homogeneous environments, the question why - and how - heterogeneous structure emerges is intriguing. We find a novel function of self-organisation in biology at the colony margin of the fungus *Aspergillus niger*. Here, the collection of interconnected hyphae splits into two subpopulations with differential expression of amylolytic enzymes. These enzymes are secreted and together break down starch in a multi-step process.

We show with a mathematical model that concentrating production and secretion of all enzymes by part of the hyphae results in more efficient substrate degradation, as compared to equal distribution over all hyphae. This model result is corroborated by experimental observations of increased metabolic activity of the wild type that displays hyphal heterogeneity as compared to a mutant that lost this heterogeneity.

The intermediate product (iso)maltose is known to induce upregulation of amylolytic enzymes. Incorporating this in the model enables the formation of hyphal heterogeneity through self-organisation. We generalize our results to a large class of multi-step processes and demonstrate wide applicability. Importantly, this shows that self-organisation may not be incidental, but favorable and thus selectable since it enhances efficiency. Due to its impact on system functioning, its study is not only of academic interest but also of practical value.

## 1. Introduction

The mycelium of filamentous fungi consists of an interconnected network of hyphae. Individual hyphae extend at their apices by polarized growth and branch subapically [1,2]. Filamentous growth allows fungi to rapidly explore and colonize various organic substrates, such as lignocellulose and starch [3]. This makes fungi key players in global nutrient recycling, as well as notorious food spoilage agents. In addition, they are utilized as cell factories for the production of enzymes and organic acids.

Starch is the most abundant plant storage carbohydrate. For instance, many crops produce starch-storing organs, such as cereals, tubers, roots, and beans [4]. Starch granules consist of amylose, a linear glucan polymer, and amylopectin, which is a highly branched glucan polymer. Fungi secrete a variety of starch-degrading enzymes divided over three functional groups of the Glycoside Hydrolase (GH) families [5]. These enzymes hydrolyze starch synergistically to ultimately produce glucose. *α*-amylases GH13 (EC 3.2.1.1) endolytically hydrolyze *α*-1,4 linkages in amylose (*α*-1,4-glucan) chains, breaking up amylose into smaller glucose polymers and maltooligosaccharides. Glucoamylases GH15 (EC 3.2.1.3) and *α*-glucosidases GH31 (EC 3.2.1.20) exolytically hydrolyze *α*-1,4 linkages from non-reducing ends, releasing glucose of *β*-anomeric and *α*-anomeric forms, respectively. Glucoamylases also hydrolyze *α*-1,6 linkages at the branch connections in amylopectin [6]. Since we focus on amylose degradation as example, degradation of amylopectin is not described here.

In *Aspergillus*, the expression of amylolytic genes is tightly regulated. The production of amylolytic enzymes is induced by (iso)maltose through activation of the transcriptional regulator AmyR which then transits to the nucleus [7] and binds to the promoters of amylolytic genes which activate transcription [8]. Thus, three enzyme families work in concert to break down amylose polymers to their constituent glucose monomers. Subsequently, these monomers pass through sugar transporter transmembrane proteins [9] and are further catabolized. We refer to both enzymes and transporters as catalysts.

Previously, the mycelium was regarded as homogeneous, but over the past decades heterogeneity has been established at multiple levels [10–13]. Hyphae of filamentous fungi are compartmentalized by cross walls. These septa have a central pore that can be plugged by Woronin bodies [14,15]. Septal plugging allows fungi to develop hyphal heterogeneity, by containing local gene expression, while still providing control over selective transport between compartments [16,17].

This was first observed for heterogeneous expression of *glaA* (glucoamylase) in the leading hyphae of *A. niger* [11], where the environmental conditions are arguably identical [11,13,18]. Two subpopulations of hyphae, either highly or lowly expressing GFP under control of the *glaA* promoter, were identified based on fluorescence intensity. This was substantiated by the observation that additional regulatory and amylolytic enzyme encoding genes follow the same expression pattern as *glaA*, i.e. *amyR* and *aamA* (*α*-amylase), respectively [13]. Moreover, hyphae highly expressing *glaA* also contain a higher abundance of 18S rRNA, indicating a higher translational activity. Taken together, this indicates that there are at least two subpopulations of hyphae in the colony margin, having either high or low transcriptional and translational activity. Notably, despite the different behavior, both subpopulations display similar growth rates under normal conditions [13].

Currently, it is not fully clear how and why filamentous fungi develop hyphal heterogeneity [19]. Previously, this phenomenon has been attributed to stochastic processes that introduce diversity within the hyphal population [11], serving as a bet-hedging strategy to enhance survival under adverse conditions [20]. Through mathematical modeling, we demonstrate that hyphal heterogeneity significantly enhances the efficiency of enzymatic starch degradation. This model prediction is corroborated by metabolic measurements comparing wild type (heterogeneous hyphae) and mutant (homogeneous hyphae) strains: heterogeneous hyphae exhibit higher metabolic activity. Extension of our model with upregulation of amylolytic enzyme concentrations via *amyR*, induced by (iso)maltose, facilitates self-organisation of hyphal heterogeneity. This finding indicates that only minimal regulatory parameters are required to improve the efficiency of the multi-step enzymatic process.

## 2. Methods

### (a) Mathematical model

The degradation of amylose by the fungus *Aspergillus niger* can be viewed as a chain of catalysed reactions. We built a concise model in which amylose, oligosaccharides, external and internal glucose are represented by *P*_1_, *P*_2_, *P*_3_ and *P*_4_ respectively - thus avoiding having to keep track of all polysaccharide concentrations separately. The actors *A*_*i*_ that catalyse the conversion from *P*_*i*_ to *P*_*i*+1_ are *α*-amylase (*A*_1_), glucoamylase combined with *α*-glucosidase (*A*_2_) and glucose transporter (*A*_3_). The concentrations of the products *P*_*i*_ and catalysts *A*_*i*_ are denoted by *p*_*i*_ and *α*_*i*_, respectively. The amylolytic process and the mathematical model are depicted in Fig. 1. Our qualitative results do not depend on the choice of model parameters unless stated otherwise. For simplicity, we set all parameters equal to 1 (our qualitative results will not depend on their values), except for the catalyst concentrations *α*_*i*_ which are allowed to vary between 0 and 2. The model equations and sensitivity analysis can be found in the supplementary material, with in Fig. 5A a more extensive model including the regulatory network.

**Figure 1.**
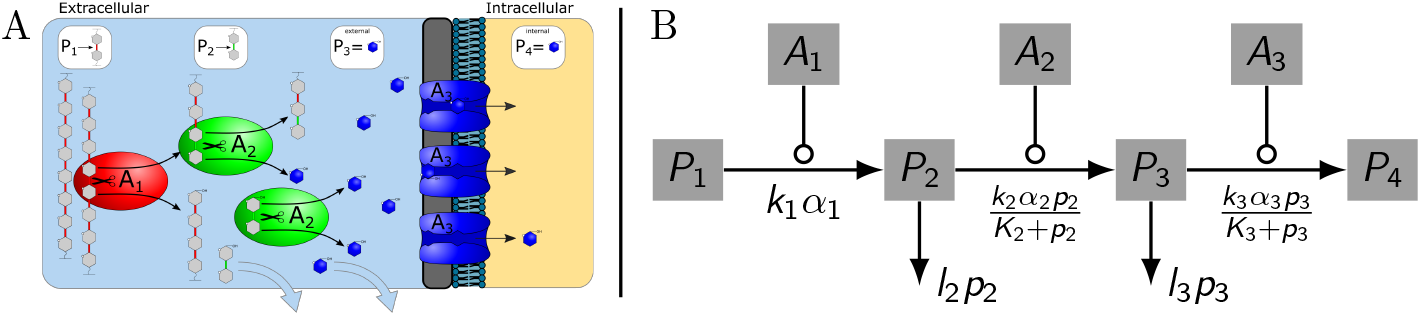
(A) Amylose degradation by *A. niger*. The *α*-amylase (A1) dissects amylose by removing a bond between units of glucose (P1). This results in one extra molecule having a bond connected to a non-reducing end (P2), which can now be dissected by the enzymes *α*-glucosidase and glucoamylase (A2) yielding external glucose monomers (P3). These monomers are transported (through A3) into the hypha (P4). Oligosaccharides (represented by P2) and external glucose are lost at a fixed rate through diffusion or uptake by rival organisms. (B) Mathematical model. A sequence of products *P*_*i*_ (concentrations *p*_*i*_) undergoing reactions catalysed by *A*_*i*_ (concentrations *α*_*i*_). *P*_1_ is assumed to be present in high concentration so that the reaction rate towards *P*_2_, *k*_1_ *α*_1_, does not depend on *p*_1_. Other reactions are according to Michaelis-Menten [21] which is the most fundamental model for enzyme kinetics: 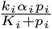. For facilitated transport this form holds under the assumption that the concentration *p*_4_ is (kept) low, so backward transport is negligible [22]. For glucose uptake by *A. niger* the Michaelis-Menten kinetics was confirmed experimentally [23]. Intermediates *P*_2_ and *P*_3_ undergo a loss of *l*_2_ *p*_2_ and *l*_3_ *p*_3_, respectively.

### (b) Strains and culture conditions

*Aspergillus niger* N402 was used as a control strain, exhibiting heterogeneous gene expression. The strain N402 *ΔhexA* with an inactivation of *hexA*, resulting in the elimination of Woronin bodies, was used as a strain exhibiting no heterogeneous gene expression [20].

To obtain spores, both strains were grown for7days on complete medium (minimal medium (MM) [24] with 0.2% tryptone, 0.1% casamino acids, 0.1% yeast extract, 0.05% yeast ribonucleic acids) supplemented with 1% glucose. Spores were harvested in saline tween (ST) (0.9% NaCl (w/v), 0.005% Tween-20 (v/v)) and counted with a haemocytometer and further diluted to a concentration of 5×10^5^ sporesml^*−*1^ in ST.

The strains were grown as sandwich colonies [10] to prevent sporulation and to allow transfer to inducing medium. The sandwich colonies were placed on top of MM agar medium supplemented with 200 mM xylose in a 9cm Petri dish, to repress amylolytic gene expression. To this end, one membrane (polycarbonate membrane, diameter 76mm, pore sixe 0.1 *µ*m; Osmonics, GE Water Technologies, Trevose, PA, USA) was placed on the medium. Thereafter, a 20 *µ*l drop of 1.25% agarose was placed on the center of the membrane. After solidification, 2 *µ*l of the spore solution (5×10^5^ sporesml^*−*1^) was placed on the agarose and allowed to dry and germinate overnight at 30°C. Then, the second membrane was placed. Sandwich colonies were incubated at 30°C for an additional4days. Subsequently, the sandwich colonies were transferred onto MM agar medium supplemented with 2% starch, to induce amylolytic activity. The colonies were placed back at 30°C for 8 or 24 hours. After the removal of the sandwich colonies, Lugol staining was performed with 10 ml Lugol solution for10min (Iodine/Potassium iodide solution, Fluka) to detect starch. Subsequently, the plates were washed three times with10 ml phosphate-buffered saline (PBS) to remove excess staining solution.

### (c) CO_2_ and biomass measurements

Carbon dioxide production for each colony was measured every 20 seconds during3minutes after amylolytic genes had been induced for 8 h on starch. The measurements were carried out in septuplicate. To this end, one culture was placed at a time in a clear airtight box (Modula 2000 ml, Mepal) together with a battery powered bespoke CO_2_ data logger inside. The box was closed to allow CO_2_ building up. The data logger was produced by connecting a SCD41 CO_2_ sensor to an Arduino Nano V3 via I2C. A SSD1306 0.96 inch oled IIC Serial White OLED Display Module (128 × 64 pixels) was also connected through I2C to output the sensor readings every 5 seconds. A bespoke PCB was designed in Fritzing to connect all hardware by soldering. The device was powered through the ISP header pins Vcc and Gnd via a 3 AA cell battery holder. The Arduino code to drive the logger and the PCB design are available on request.

The fungal biomass was quantified by harvesting the sandwich colony. To this end, the sandwich colony was placed in a new petri dish and the top membrane was removed. The colony was flipped over and the second membrane was removed. The mycelium was subsequently dried for 5 days at 60°C and the resulting dry weight of the biomass was weighed.

### (d) Statistics

The differences in metabolic activity between N402 and N402*ΔhexA* were assessed by dividing the absolute difference in carbon dioxide levels over3minutes (ppm) by the total biomass (mg) of each colony. The assumption of normality was tested for each data set using a Levene’s test. The p-value and confidence interval were tested using a Two Sample t-test. The power was tested using Vanderbilt PS: Power and Sample Size Calculation version 3.1.6, October 2018 (Dupont & Plummer, n.d.).

## 3. Results

### (a) Model: heterogeneous is more efficient

We use a mathematical model (see section2(a)) to study how concentrating catalysts at hyphae influences the efficiency of starch degradation. For each distribution of the catalyst concentrations *α*_1_ (*α*-amylases), *α*_2_ (glucoamylases combined with *α*-glucosidases) and *α*_3_ (glucose transporters) we determined the speed at which hyphae absorb glucose - the harvest rate - in equilibrium. For the sake of simplicity, we restrict our attention to two otherwise identical hyphae, and vary the distribution of the catalysts under the constraint that the total amount of each catalyst remains constant.

First suppose that at both hyphae *α*_*i*_ = 1 for each catalyst so that both hyphae harvest the same amount of glucose. This is the homogeneous benchmark (Fig. 2; orange scenario). Now, if instead the first catalyst is concentrated at one hypha, the other hypha stops harvesting. In case the other catalysts are evenly split, this results in a lower average harvest rate than the benchmark (Fig. 2; red scenario). With all three catalysts concentrated at one hypha, the amylolytic process is more efficient than the benchmark (Fig. 2; blue scenario). Conceptually, only concentrating the first catalyst leads to congestion at subsequent steps and relatively high loss terms, while concentrating all catalysts yields high throughput and relatively low losses.

**Figure 2.**
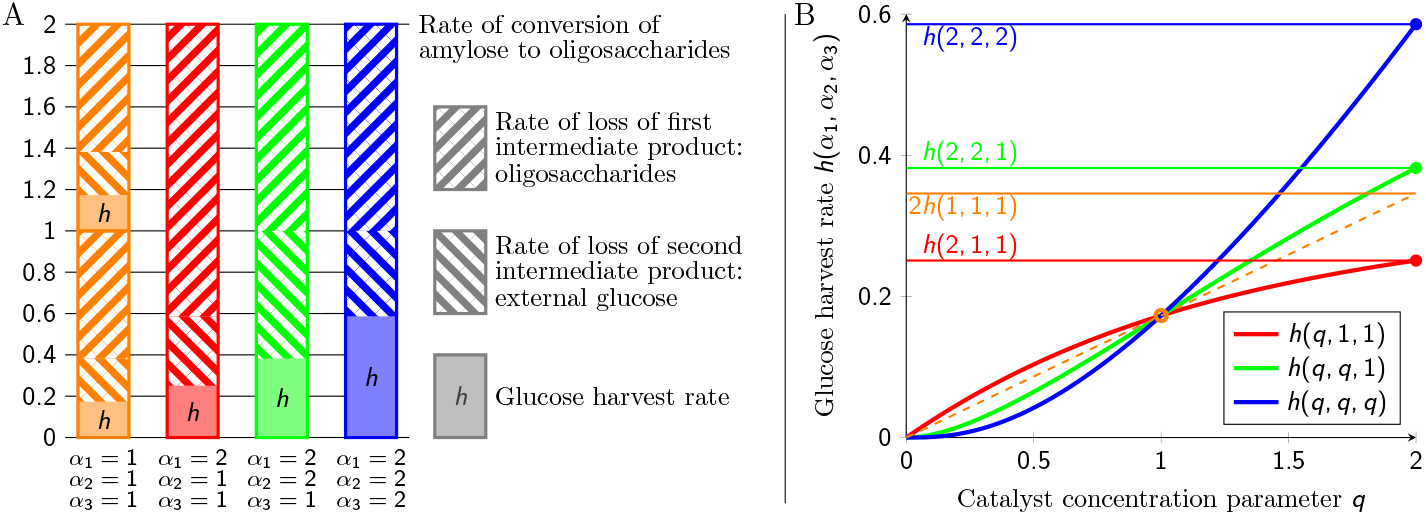
The process of amylose degradation depending on how catalyst is split over two hyphae, keeping the average catalyst concentrations *α*_1_, *α*_2_, *α*_3_ equal to1. (A) The total rate of conversion of amylose, the first step in the three-step process, is the same in all scenarios and equals2. Intermediates (oligosaccharides and external glucose) are lost due to diffusion or uptake by rival organisms, resulting in a lower glucose harvest rate, which is dependent on the scenario. Orange scenario (benchmark): fifty-fifty distribution of all catalysts, so that *α*_1_ =*α*_2_ =*α*_3_ = 1in both hyphae, resulting in a harvest rate of *h* (1,1,1) *≈* 0.17 twice. Red: only *α*_1_ = 2 is concentrated at one hypha; *α*_2_ and *α*_3_ are evenly split. Harvest is restricted to one hypha with rate *h*(2,1,1) *≈* 0.25 *<* 2*h*(1,1,1), so smaller than the benchmark. Green: additionally *α*_2_ is concentrated at one hypha; *α*_3_ remains evenly split (*α*_3_ = 1). Harvest is restricted to one hypha with rate *h*(2,2,1) *≈* 0.38 *>* 2 *h*(1,1,1), so larger than the benchmark. Compared to the red scenario, concentrating *α*_2_ results in a drop of the loss of the first intermediate product, but the loss of the second intermediate increases. Blue: also *α*_3_ is concentrated at one hypha. Harvest is restricted to one hypha with rate *h*(2,2,2) *≈* 0.59 which outperforms all other scenarios. (B) Harvest rate as a function of catalyst concentrations for the three heterogeneous scenarios. All curves pass through the benchmark (orange circle) at *q* = 1. Since the red curve has a decreasing slope, concentrating only *α*_1_ yields a diminishing return of investment. The blue curve has an increasing slope, so there is an increasing return of investment: economy of scale. The green curve is sigmoidal and because here at *q* = 1 the slope is increasing, it also has an increasing return of investment. Explicit formulas for the curves can be found in the supplementary material.

The intermediate scenario is where two out of three catalysts are concentrated at one hypha (Fig. 2; green scenario). Concentrating the second catalyst *α*_2_ results in less congestion at the second step, but partially at the expense of more congestion at the third step. In Fig. 2 the harvest rate is higher than that of the benchmark. By changing the reaction constant of the first reaction *k*_1_ from the default value *k*_1_ = 1 to *k*_1_ = 10, leading to more congestion, concentrating two catalysts can also result in a harvest rate that is lower than the benchmark (supplementary material, Fig. S5).

The graphs in Fig. 2 help explain these results. The red curve has a decreasing slope, so concentrating one catalyst leads to a diminishing return of investment. So there is no incentive to concentrate only the first catalyst. The blue curve has an increasing slope, so concentrating all catalysts leads to an increasing return of investment. This economy of scale is an incentive to concentrate all catalyst at one hypha. The green curve is sigmoidal, so concentrating part of the catalysts can either lead to a diminishing or increasing return. Thus, the model shows that heterogeneity permits a higher glucose harvest rate.

Importantly, the appearance of diminishing or increasing return of investment only depends on the number of concentrating catalysts and does not depend on the choice of any parameter value. Namely, within our model framework, concentrating one catalyst always results in a diminishing return of investment. Increasing all catalyst concentrations always results in an increasing return of investment. Increasing multiple but not all catalyst concentrations always results in a sigmoidal curve. Only the location of the switch from increasing to diminishing return depends on parameter values. The general picture is sketched in Fig. 3. Sensitivity analysis, generalisation and mathematical proofs can be found in the supplementary material.

**Figure 3.**
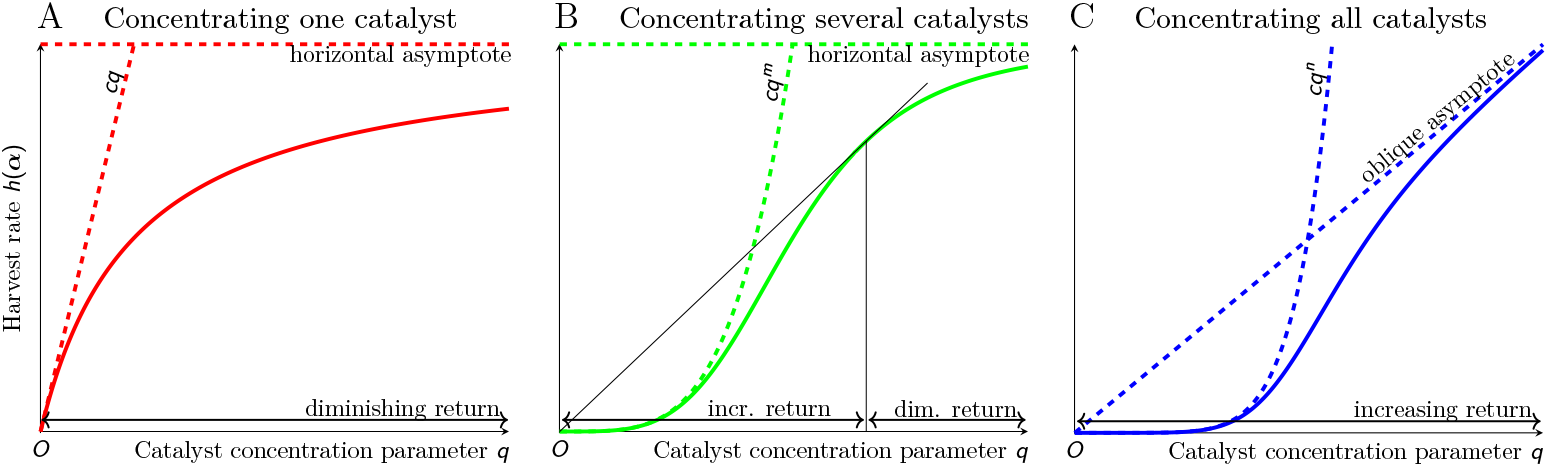
General picture of the dependence of the harvest rate on the catalyst concentration parameter *q*, depending on the number of catalyst concentrations that depend on *q*, which is independent of the value of all other parameters. The limiting behavior for low and high *q* is determined by Corollaries 3 and 4 of the supplementary material. (A) Only one catalyst concentration depends on *q*. In this case, there is always a diminishing return of investment (supplementary material, Corollary 1). (B) Multiple *m* (but not all *n*) catalyst concentrations depend on *q*. The harvest rate is a sigmoidal function so that concentrating multiple (but not all) catalysts can both lead to an increasing and diminishing return of investment (supplementary material, Corollary 5). See supplementary material Fig. S4 and S5 where this is worked out for the amylolytic model. (C) All *n* catalyst concentrations depend on *q*. In this case, there is always an increasing return of investment (supplementary material, Corollary 2).

## (b) Experimental validation

Both the amylolytic regulatory gene *amyR* and the amylolytic enzymes encoding genes *aamA* (*α*-amylase, *α*_1_) and *glaA* (glucoamylase, *α*_2_), are heterogeneously co-expressed in the wild type [11,13]. The absence of Woronin bodies results in the elimination of septal plugging and thereby the loss of heterogeneity of amylolytic gene expression in the *hexA* mutant [16,20]. We used these strains to test the outcome of our model: that concentrating catalysts permits a higher glucose harvest rates than homogeneously distributing all catalysts.

To this end, sandwich colonies of the *A. niger* strains N402 (wild type) and N402*ΔhexA* (mutant) were grown for 5 days on 200 mM xylose, after which they were transferred for8hours to MM agar plates supplemented with 2% starch to induce amylolytic activity [25]. After 8 hours starch was still present underneath the colonies, while most starch was degraded after 24 hours (supplementary material, Fig. S7). Thus, the metabolic activity of both strains was determined after8hours by measuring the carbon dioxide production over 3 minutes, corrected for biomass.

A significant difference in metabolic activity between the wild type and the mutant strain was observed. On average, the metabolic activity (CO_2_ production/mg biomass) was 48% higher (*p* = 0.043, power = 1.00) in the heterogeneous wild type with respect to the mutant strain. This implies that also the glucose harvest rate of the wild type is higher than that of the mutant strain. But there was no apparent distinction between the amount of remaining amylose (starch) for the two strains after 8 and 24 hours (supplementary material, Fig. S7). Of note, lugol can only bind the helix structure of starch, but not its degradation products, which lack this structure. The difference in harvest rate is therefore not due to a difference in conversion rate from amylose to oligosaccharides, but must be due to the efficiency of heterogeneity in subsequent conversion steps, which is in agreement with the model (Fig. 2).

In contrast, the average biomass of N402 and N402*ΔhexA*, 69.31 and 68.01 mg, respectively, did not show a significant difference (*p* = 0.58, power = 0.07). Most of the growth occurs during the preliminary 5 days on xylose. Since uptake of xylose is a one-step process, here cellular heterogeneity is not predicted to have an effect according to our model, which explains why the average biomass is not significantly different. Thus, the biological experiments support the model findings.

### (c) Model: self-organisation enables heterogeneity

In the previous sections we have shown that starch degradation by *A. niger* is more efficient if the hyphae specialise into subpopulations of highly active and lowly active hyphae. We now sketch how, with minimal adaptation of the model, this heterogeneity can be realized through self-organisation.

It is known that the production of amylolytic enzymes (with concentration *α*_1_ and *α*_2_) is induced by (iso)maltose (represented by concentration *p*_2_) by upregulation through AmyR [7]. Since increasing *α*_1_ results in a higher production of *p*_2_, this may yield a positive feedback.

The induction by (iso)maltose depends on the external concentration of maltose and the amount of maltose transporters. Maltose itself leads to increased expression of maltose transporters ([26] and reviewed by [27]). Moreover,*α*-gluscosidases in *Aspergillus* catalyse the formation of isomaltose [25]. For these reasons, the strength of the induction is likely a non-linear function of external maltose concentration *p*_2_. We postulate that *α*_1_ and *α*_2_ both on *p*_2_ through a function that resembles a sigmoid (also found in [28]). For the sake of simplicity we choose 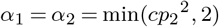. This means that they depend quadratically on *p*_2_ but cannot increase beyond the value 2.

The coefficient *c* controls the strength of the upregulation. Provided *c* is sufficiently large, this results in bi-stability of highly and lowly active hyphae. In Fig. 4, three bi-stable configurations are depicted. Each hypha can only reside in one of the two stable equilibria (*p*_2_ = *α*_1_ = *α*_2_ = 0 or *p*_2_ = 1and *α*_1_ = *α*_2_ = 2) corresponding to low or high activity.

**Figure 4.**
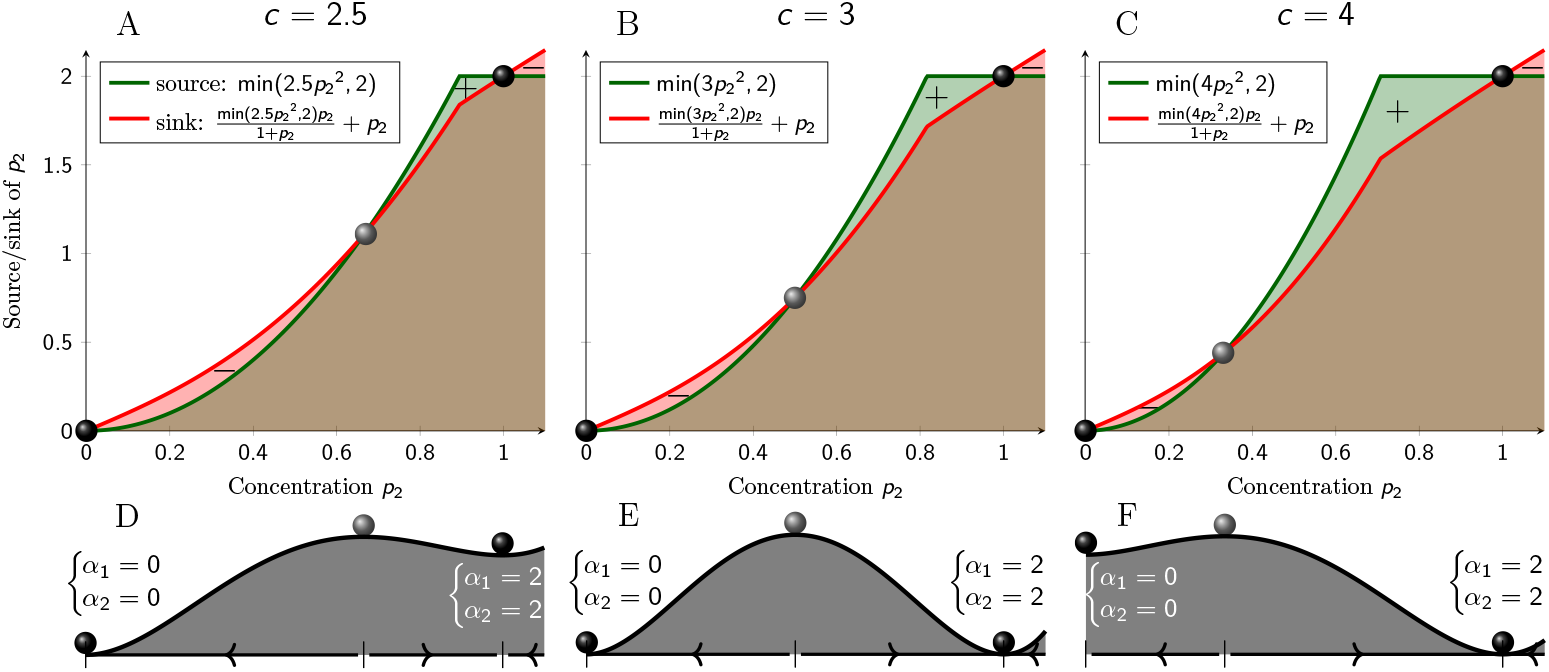
(A-C) Source term (green curve) and sum of two sink terms (red) of *p*_2_ (related to oligosaccharides) as a function of *p*_2_ itself, with upregulation 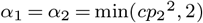 with varying strength: *c* = 2.5 (A), *c* = 3 (B) and *c* = 4 (C). Equilibria when source and sink terms are equal are depicted by balls, which are either stable (black) or unstable (gray). (D-F) The coefficient *c* modulates the location of the unstable equilibrium in the middle and thus the basins of attraction of the two stable equilibria, where either *α*_1_ = *α*_2_ = 0 (on the left) or *α*_1_ = *α*_2_ = 2 (on the right), for *c* = 2.5 (D), *c* = 3 (E) and *c* = 4 (F).

The strength of the upregulation *c* determines how the basins of attraction (bounded by the unstable equilibrium) are positioned. A larger basin of attraction generally leads to a larger proportion of hyphae converging to the corresponding equilibrium. By modulating the percentage of highly active hyphae the glucose concentration inside the mycelium may be managed. Too many active hyphae would result in a high internal glucose concentration, which induces the retention of CreA to the nucleus, which in turn inhibits the production of amylolytic enzymes [29]. The strength of the upregulation *c* thus decreases and so would the percentage of highly active hyphae, reducing the influx of glucose. This way, the hyphae would dynamically self-organize into a lowly and highly active subpopulation. Varying proportions between lowly and highly active hyphae of *A. niger* have been observed, depending on the conditions [11].

It is currently not known whether the concentration of glucose transporters *α*_3_ in the hyphal cell membrane also depends on *p*_2_ like *α*_1_ and *α*_2_. For bi-stability of the system this is irrelevant, because *α*_3_ does not influence *p*_2_. However, it would improve the efficiency, namely from the green scenario in Fig. 2 to the blue scenario. The dependence of *α*_3_ on *p*_2_ could also be indirect, e.g. through dependence of *α*_3_ on the external glucose concentration *p*_3_ [30]. Either way, we conclude that a minimal model adjustment enables heterogeneity through self-organisation, and more efficient starch degradation.

## 4. Discussion

In this study we showed that self-organisation optimizes multi-step processes. Specifically, we showed by both modeling and biological experiments that hyphal heterogeneity in *A. niger*, observed in [11], enhances the efficiency of starch degradation. For simplicity, we restricted ourselves to the linear amylose component of starch in the model but it is extendable to amylopectin by including a preliminary reaction step. Within our general multi-step mathematical framework we prove that concentrating one catalyst is never beneficial (Fig. 3). Only if multiple (preferably all) catalysts are concentrated this permits a higher overall harvest rate. Master controller genes or transcription factors such as AmyR affecting multiple steps at once are thus not only convenient but also necessary for evolution towards more efficient heterogeneous transcription.

Our model analysis assumes that the glucose harvest rate is in equilibrium. After exposure of *A. niger* to starch and induction of amylolytic enzymes, hyphae with lower catalyst concentrations need longer to reach their equilibrium harvest rate, because the concentration of intermediates in this case builds up relatively slowly. Part of the difference in metabolic activity observed in our experiment may be due to the *hexA* mutant not having reached its equilibrium harvest rate, but this only underscores the efficiency of heterogeneity.

Our results relate to the concept of *dissipative structure* coined by Ilya Prigogine (Nobel prize in chemistry, 1977). Here, “dissipative” refers to a thermodynamically open system, which need not evolve to equilibrium. Because there is a continuous supply of amylose, the amylolytic model is also open and the differences in activity of the hyphae is linked to dissipative structure. In [31] it was stated that “biological structures can only originate (…) and be maintained by a continuous supply of energy”. Here, it was also contemplated that “In the inhomogeneous state (…) the overall reaction rate is increased”. In this article, we indeed have shown how, within our multi-step model framework, heterogeneity permits a higher over-all reaction rate. This is at odds with the previous perception that heterogeneity leads to sub-optimal production in e.g. bioreactors, since not all micro colonies participate in production [18,32]. Instead, we find that the maximum production level can only be attained through heterogeneity. Thus, contrary to what the name might suggest, dissipative structure is economic.

For our general model framework, the qualitative result that concentrating multiple catalysts leads to economy of scale holds for a wide range of parameter values (Fig. 3B). Concentrating all catalysts always leads to economy of scale (Fig. 3C), so independent of all parameter values. By natural selection of efficient processes we thus expect that our results may be applicable in other kingdoms of life at possibly other length scales. Necessary and sufficient conditions within our framework are 1) a multi-step process where the reaction speed of multiple steps can be influenced, 2) sufficient separation to allow differentiation (in case of amylolysis by septal plugging impeding cytoplasmic mixing) and 3) some loss of intermediate products. In amylolysis, the intermediates are external glucose and oligosaccharides. As long as the intermediates have not been transported into the hyphae, there is loss due to diffusion or uptake by rival organisms. With the three conditions in mind, we end by highlighting similarities, applicability, and limitations of the model framework to four more cases in a broad range of contexts (Fig. 5B-E).

**Figure 5.**
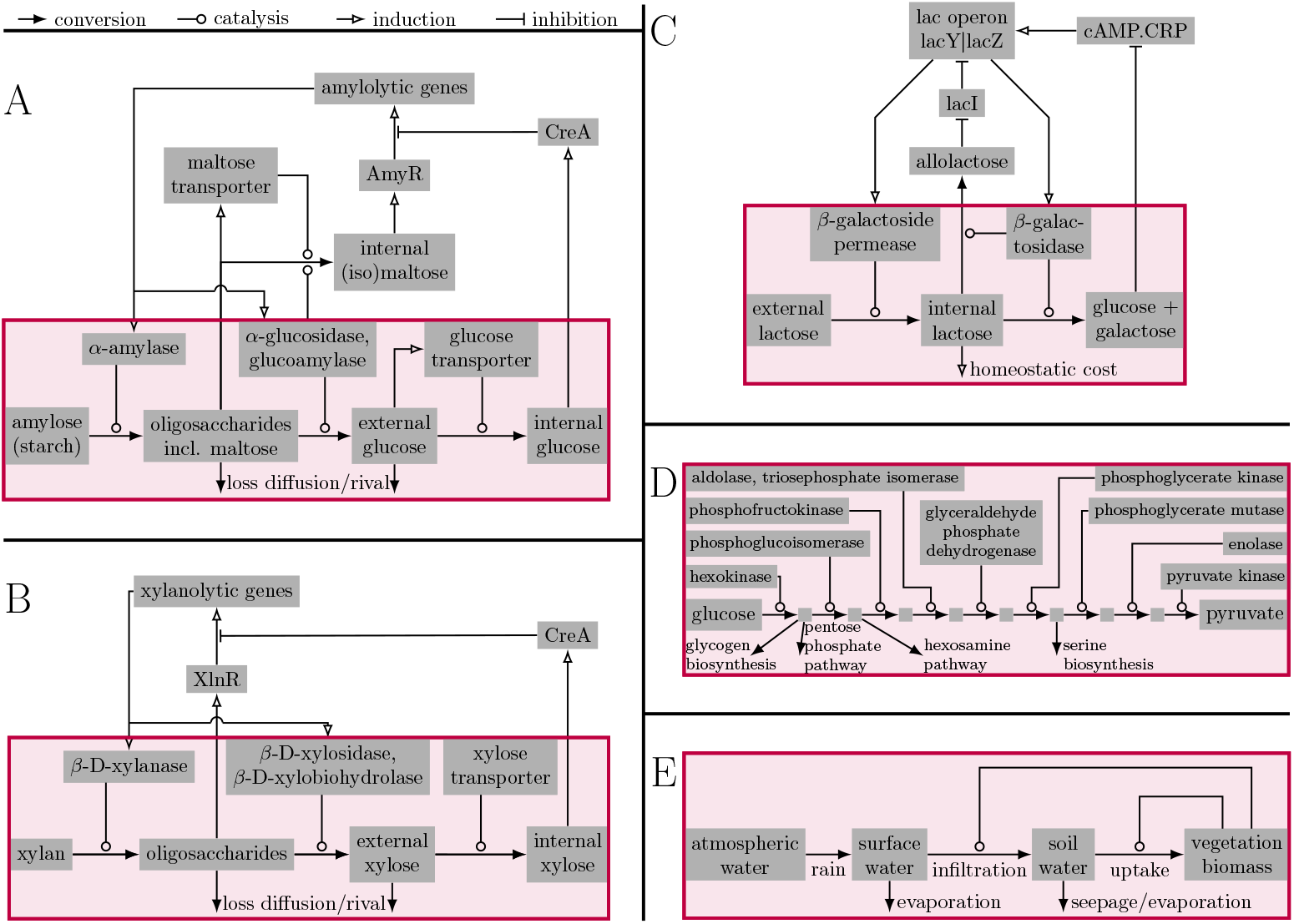
Similarity between A) amylolysis and B) xylanolysis by *A. niger*, C) uptake and digestion of lactose by *E. coli*, D) glycolysis and E) water harvesting by dryland vegetation. A) Amylolytic pathway. Within the box the minimal model of catalysed degradation of starch, with regulatory network outside the box. (Iso)maltose promotes transcription of amylolytic genes through upregulation of AmyR [7], resulting in production of amylolytic genes. Maltose leads to increased expression of maltose transporters ([26] and reviewed by [27]), and *α*-gluscosidases in *Aspergillus* catalyse the formation of isomaltose [25]. The external glucose concentration influences the presence of glucose transporters [30]. The end product, internal glucose, inhibits the production of enzymes through the catabolic respressor CreA [29]. B) Xynalolytic pathway. The minimal model (within red box) and regulatory network is very similar to that of amylolysis. XlnR regulates the expression of xylanolytic enzymes [33], which break down xylan sequentially (reviewed in [34]). Xylooligosaccharides trigger the expression of XlnR [35], transcription is again down-regulated by CreA [36]. C) Lactolytic pathway. The red box shows a minimal model. Both the membrane protein that enables transport of lactose into the cell and the enzyme that cleaves lactose are induced by the intermediate (allo)lactose [37], which is similar to A) and B), but this time through inhibition of an inhibitor. Also, glucose again inhibits the production of the catalysts: here glucose represses cAMP, which after binding to its receptor protein (CRP) would promote the production of catalyst [38]. Since the intermediate - internal lactose - is intracellular, it may not be lost through diffusion or uptake by rival organisms. Loss may be interpreted more broadly by acknowledging that maintaining an intracellular metabolite concentration induces a homeostatic cost. D) Glycolytic pathway. Only the minimal model without regulatory network is shown. Ten enzymes together sequentially convert glucose into pyruvate. The reactions are again intracellular, loss of intermediates may in this case be interpreted by other pathways competing for intermediates. E) Harvesting of rain water by dryland vegetation. Here the end product vegetation itself acts as a catalyst, increasing both the infiltration and the uptake rate. This is opposite to A-C, where end products inhibit the production of catalysts (to prevent overproduction). In the dryland context there is no regulatory network.

Starting close to home, next to starch, *Aspergillus* can degrade many more complex substrates, and this degradation is generally mediated by specific transcriptional regulators such as AmyR. For example, XlnR regulates the expression of xylanolytic enzymes [33], which break down xylan sequentially (reviewed in [34]), see Fig. 5B. Xylooligosaccharides trigger the expression of XlnR [35], transcription is again down-regulated by CreA [36]. As degradation of xylan proceeds in multiple steps, concentrating enzyme production in part of the hyphae again permits a more efficient substrate degradation. Expression of xylanolytic genes is known to be heterogeneous: the production of *faeA* and *aguA* is again separated in two distinct hyphal populations [13]. The corresponding enzymes remove side chains, which could be included in the model as a preliminary step. Whether the enzymes that degrade the main chain are also heterogeneously expressed remains to be investigated.

Secondly, the lac operon of *E. coli* [37] regulates two steps in the digestion and uptake of lactose (Fig. 5C). Specifically, both the membrane protein that enables transport of lactose into the cell and the enzyme that cleaves lactose are induced by (allo)lactose (condition (1)). Since the intermediate - internal lactose - is intracellular, it cannot be lost through diffusion or uptake by rival organisms. Loss may be interpreted more broadly by acknowledging that maintaining an intracellular metabolite concentration incurs a homeostatic cost (condition 3). The overall homeostatic cost at the population level would then be lower if cleavage of lactose is performed in only part of the bacteria (keeping overall catalyst concentration constant), while the harvest rate remains the same, thereby potentially increasing overall efficiency. On the population level, there is sufficient separation for heterogeneity (condition (2)). Leakage [39] of glucose or galactose could be part of a division of labour; specialisation by bacteria has been observed in biofilms [40]. The lac operon was found to be bi-stable for induction by artificial inducer but less likely so for the natural inducer (allo)lactose [41–43].

In the previously discussed cases, sufficient separation for differentiation (condition (2)) was ensured because hyphae or entire cells acted as compartments. On a smaller scale, subcellular compartmentalisation facilitates metabolite or substrate channelling: the transfer of intermediates from one enzyme to the next [44–46]. A prominent example of channelling is glycolysis (reviewed in [47]). Here, ten enzymes catalyse the conversion of glucose into pyruvate (condition (1)), see Fig. 5D. Intermediates of glycolysis are involved in side reactions, which are then lost from the glycolytic pathway (condition (3)). Channelling is a way to regulate the metabolic flux network and improve the efficiency of the channelled pathway by lowering the chance that intermediates are lost from that pathway [48–50] or to protect homeostasis by keeping the concentration of intermediates low [51]. The end product of glycolysis, pyruvate, is subjected to oxidative decarboxylation by the pyruvate dehydrogenase complex of three enzymes, which is also a form of substrate channelling. The resulting acetyl-CoA enters the Krebs cycle, with corresponding enzymes concentrated in mitochondria, another from of substrate channelling by subcellular localization. The entire chain from glycolysis up to and including the Krebs cycle could function as one partially channelled pathway: if the respiration rate increases, glycolytic enzymes dynamically associate to the outer surface of mitochondria [52]. Other examples of dynamic substrate channelling are hypoxic stress leading to compartmentalisation of glycolysis enzymes [53] and the level of purine influencing the association and dissociation of enzyme clusters in purine biosynthesis [54].

As a fourth and final case, we apply our model framework to vegetation patterns in drylands. These patterns were first reported in 1950 based on air photography [55]. With distances of tens to hundreds of meters between patches of vegetation, interspersed with bare soil, this heterogeneity operates on an entirely different scale. An explanation for the emergence of self-organisation is that the water infiltration rate is higher within vegetated patches [56,57], so there is a local facilitation effect, similar to biofilms. In a second step, infiltrated water is taken up by the vegetation. The reaction speed of both steps, infiltration and uptake, is increased by vegetation (condition (1), Fig. 5E). Vegetation can separate into patches over generations (condition (2)). And if water is not taken up, it evaporates or seeps away (condition (3)). Models for dryland vegetation patterns that incorporate the infiltration feedback, e.g. [58,59], fit our model framework. Accordingly, clustered vegetation harvests water more efficiently than homogenously distributed vegetation, thereby improving resilience against drought (reviewed by [60]).

Dryland vegetation forms a regular Turing pattern [61,62]. A topic of ongoing research is whether the population of highly active hyphae in *A. niger* also forms a regular pattern. In our model we focused on two hyphae, just like Turing started explaining his seminal idea by considering two neighboring cells [61]. After inclusion of exchange of substances between hyphae in the model, we would expect that hyphae neighbouring highly active hyphae may also have a tendency to become highly active, resulting in clumps of hyphae that jointly break down the substrate. The size of these clumps would be determined by the spreading speed of inducing oligosaccharides like maltose in relation to that of glucose - which spreads faster and leads to catabolic repression through CreA - and the need for glucose. Around these clumps there may be hyphae that only express increased concentration of glucose transporters, harvesting external glucose without taking part in the enzymatic degradation. The resulting high internal glucose concentration would oppose the production of amylolytic enzymes in these hyphae. The spacing between clumps is arguably dependent on the mobility of glucose. If glucose moves relatively fast (e.g. via cytoplasmic streaming [63]), clumps of highly active hyphae may lie further apart from each other, since lowly active hyphae need not be in the direct vicinity of highly active hyphae to acquire sufficient glucose for homeostasis. As substrate is locally depleted, the positions of active clumps may change dynamically.

Dryland vegetation pattern formation occurs without extensive regulatory network (Fig. 5E), so the patterns are truly *self*-organised. For the metabolism of *Aspergillus*, self-organisation is facilitated by a regulatory network, so this is a type of *organised self-organisation*. In Fig. 5D, no regularity network of the glucolytic pathway is shown but it easily outcompetes the other examples in terms of complexity. Here, self-organisation may still play a role in self-assembly of compartmentalisation [47,64], but the entire process seems to be more on the organised side rather than self-organised side.

Life has evolved to carry out complex processes efficiently. By organisation and specialisation, the overall efficiency can be improved. We have shown this also applies to self-organisation in the context of multi-step processes, so higher yield is possible with minimal control. This improves fitness in situations where full control is not possible, e.g. if part of the process is extracellular. An additional benefit is that self-organised patterns can more easily adapt to changing conditions, allowing for greater flexibility as compared to more rigid structures in life. The extent to which self-organisation is employed in all kingdoms of life remains to be investigated.

## Acknowledgements

The initial phase of this research was supported by NWO TTW grant ‘Traffic control’ [15493]. R.B. was supported by NWO TTW Vidi grant ‘Feed me’ [18920]. The funders were not involved in study design, in the analysis, the writing of the article or the decision to submit for publication.

