## Supplementary for "Employment of self-organisation to achieve economy of scale in biology"

### Supplementary Material

#### Derivation and generalisation of the mathematical model for amylolysis

We first derive the model for the digestion of amylose by *A. niger*. This process consists of three steps. We then generalize the model to an arbitrary number of reaction steps and present formulas for the equilibrium harvest rate.

The three steps of amylolysis are:

- The enzyme $\alpha$-amylase, denoted $A_1$, catalyses hydrolisation of bonds in between units of glucose ($P_1$) within amylose. With each of these reactions one of these bonds is removed and a next bond becomes an additional bond neighboring a non-reducing end ($P_2$). Assuming superfluous availability of $P_1$ (and water), the corresponding reaction is $2\,\mathrm{P}_1 + \mathrm{H_2O} \xrightarrow{k_1\alpha_1} \mathrm{P}_2$.
- The enzymes glucoamylase and $\alpha$-glucosidase, together denoted $A_2$, catalyse hydrolysis of the bonds connected to a non-reducing end. Here, there a two possibilities. If $A_2$ acts on maltose (chain of two glucose units) then $P_2$ is replaced by two glucose molecules ($P_3$): $\mathrm{P}_2 + \mathrm{H_2O} \xrightarrow{\frac{k_2\alpha_2 p_2}{K_2+p_2}} 2\,\mathrm{P}_3$. If $A_2$ acts on a longer chain of glucose units, then the net effect is that the amount of $P_2$ remains unchanged and only one glucose molecule is created: $\mathrm{P}_2 + \mathrm{H_2O} + \mathrm{P}_1 \xrightarrow{\frac{k_4\alpha_2 p_2}{K_2+p_2}} \mathrm{P}_3 + \mathrm{P}_2$. In both cases, we use Michaelis-Menten (1) for the reaction speed.
- As a third step, we view the uptake of glucose by fungal hyphae facilitated by transporters ($A_3$) resulting in internal glucose ($P_4$): $\mathrm{P}_3 \xrightarrow{\frac{k_3\alpha_3 p_3}{K_3+p_3}} \mathrm{P}_4$. This is a passive process, so glucose can in theory also move in the opposite direction, but we assume this is negligible because internal glucose concentrations are kept low. In this case the rate of uptake is of Michaelis-Menten form (2), which was also confirmed experimentally (3).

Together, this yields the following mathematical model:

$$\begin{cases} \dfrac{dp_2}{dt} = k_1\alpha_1 - \dfrac{k_2\alpha_2 p_2}{K_2+p_2} - l_2 p_2 \\ \dfrac{dp_3}{dt} = \dfrac{2k_2+k_4}{K_2+p_2}\alpha_2 p_2 - \dfrac{k_3\alpha_3 p_3}{K_3+p_3} - l_3 p_3 \end{cases}. \quad [1]$$

Note that we do not keep count of $P_1$ (bonds between glucose units within amylose) since it assumed to be superfluously available. And we do not keep count of $P_4$ (internal glucose) since we are mainly interested in the harvest rate at which glucose is transported into the hypha, for which we assume that $p_4$ is kept low. After a rescaling ($k_2 p_3 = \tilde{p}_3(2k_2+k_4)$, $k_2 k_3 = \tilde{k}_3(2k_2+k_4)$, $k_2 K_3 = \tilde{K}_3(2k_2+k_4)$ and removing the tildes again) we obtain the equivalent model

$$\begin{cases} \dfrac{dp_2}{dt} = k_1\alpha_1 - \dfrac{k_2\alpha_2 p_2}{K_2+p_2} - l_2 p_2 \\ \dfrac{dp_3}{dt} = \dfrac{k_2\alpha_2 p_2}{K_2+p_2} - \dfrac{k_3\alpha_3 p_3}{K_3+p_3} - l_3 p_3 \end{cases} \quad [2]$$

which is the amylolytic model we used in the article. Our qualitative results also apply to other multi-step processes, for $n \geq 2$ steps the generalisation is as follows:

$$\begin{cases} \dfrac{dp_2}{dt} = k_1\alpha_1 - \dfrac{k_2\alpha_2 p_2}{K_2+p_2} - l_2 p_2 \\ \dfrac{dp_i}{dt} = \dfrac{k_{i-1}\alpha_{i-1} p_{i-1}}{K_{i-1}+p_{i-1}} - \dfrac{k_i\alpha_i p_i}{K_i+p_i} - l_i p_i \quad \text{for } 2 < i \leq n \end{cases}. \quad [3]$$

As shorthand for the conversion rates we use $r_1 = k_1\alpha_1$ for the conversion from $P_1$ to $P_2$ and for $i > 1$, $r_i = \frac{k_i\alpha_i p_i}{K_i+p_i}$ for the conversion from $P_i$ to $P_{i+1}$.

We restrict our attention to equilibria, where $p_2 = \bar{p}_2, p_3 = \bar{p}_3, ..., p_n = \bar{p}_n$ are constant in time. Without loss of generality we may assume that all loss rates $l_i$ are non-zero. To show this, suppose instead that $l_j = 0$ for some $j$. Then in the graphical representation of the multi-step process (Fig. S1) there is no loss term emanating from $P_j$. Considering that we are interested in equilibrium, there are two options. If $r_{j-1} < k_j\alpha_j$, in equilibrium the transition rate $r_j$ from $P_j \to P_{j+1}$ will equal the transition rate $r_{j-1}$ from $P_{j-1} \to P_j$. The two steps $P_{j-1} \to P_j \to P_{j+1}$ can thus be replaced by a single step $P_{j-1} \to P_{j+1}$. Otherwise, $r_{j-1} \geq k_j\alpha_j$, which means that the concentration $p_j$ of $P_j$ will build up indefinitely and the transition rate $P_j \to P_{j+1}$ will

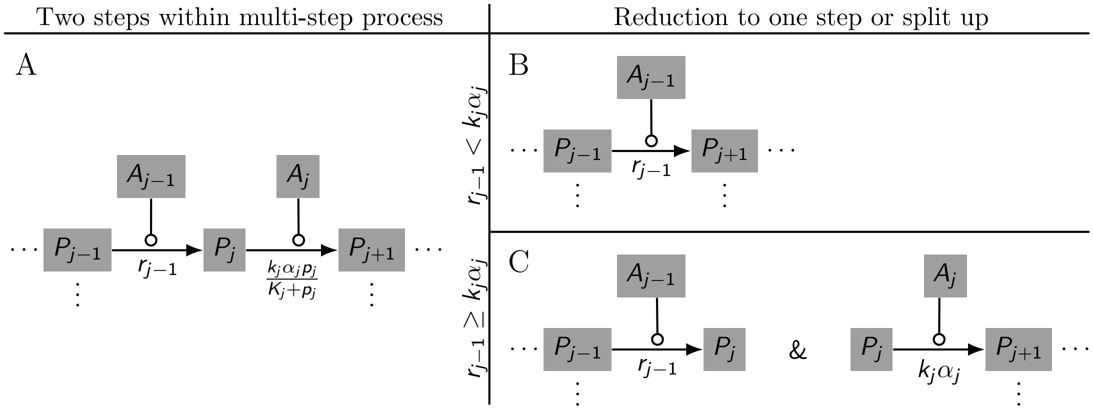


**Fig. S1.** Graphical representation of the reduction of the case $l_j = 0$. (A) Original process without loss of intermediate product $P_j$. (B) Reduction if $r_{j-1} < k_j\alpha_j$. (C) Reduction to two split processes if $r_{j-1} \geq k_j\alpha_j$.

converge to $k_j\alpha_j$. The multi-step process can thus be viewed as split up into two parts (Fig. S1). From now on we will assume that $l_i > 0$ for all $i$.

Now, for each set of fixed catalyst concentrations $\boldsymbol{\alpha} = (\alpha_1, \alpha_2, ..., \alpha_n)$ the model (Eq. (3)) has a unique equilibrium $(\bar{p}_2, \bar{p}_3, ..., \bar{p}_n)$ solving $\frac{d\bar{p}_2}{dt} = \frac{d\bar{p}_3}{dt} = ... = \frac{d\bar{p}_n}{dt} = 0$. The equilibrium is non-negative, stable and each initial condition over time converges to this equilibrium (i.e. it is a global attractor). In equilibrium, we refer to the steady production of the end product $p_{n+1}$ as the harvest rate $h(\boldsymbol{\alpha})$, which is a function of the enzyme distribution $\boldsymbol{\alpha}$.

The equilibruim concentration of each $P_i$ $(2 \leq i \leq n)$ is given by

$$\bar{p}_i(\boldsymbol{\alpha}) = \frac{1}{2l_i}\left(\beta_i - k_i\alpha_i - K_il_i + \sqrt{4\beta_iK_il_i{}^2 + (\beta_i - k_i\alpha_i - K_il_i)^2}\right) \quad [4]$$

where $\beta_2 = k_1\alpha_1$ and $\beta_i = \frac{k_{i-1}\alpha_{i-1}\bar{p}_{i-1}}{K_{i-1}+\bar{p}_{i-1}}$ for $i > 2$. The harvest rate is given by

$$h(\boldsymbol{\alpha}) = \frac{k_n\alpha_n\bar{p}_n(\boldsymbol{\alpha})}{K_n + \bar{p}_n(\boldsymbol{\alpha})} = k_1\alpha_1 - (l_2\bar{p}_2(\boldsymbol{\alpha}) + l_3\bar{p}_3(\boldsymbol{\alpha}) + ... + l_n\bar{p}_n(\boldsymbol{\alpha})). \quad [5]$$

It is possible to find $h(\boldsymbol{\alpha})$ as an explicit function of the $\alpha_i$. For the amylolytic model (Eq. (2)), with all other parameters set equal to one, it can be expressed as:

$$\begin{aligned} h(\boldsymbol{\alpha}) =& \alpha_1 - \frac{1}{2}\left(\alpha_1 - \alpha_2 - 1 + \sqrt{4\alpha_1 + (\alpha_1 - \alpha_2 - 1)^2}\right) - \frac{1}{2}\Bigg\{\alpha_2\frac{\alpha_1 - \alpha_2 - 1 + \sqrt{4\alpha_1 + (\alpha_1 - \alpha_2 - 1)^2}}{2 + \alpha_1 - \alpha_2 - 1 + \sqrt{4\alpha_1 + (\alpha_1 - \alpha_2 - 1)^2}} - \alpha_3 - 1 \\ &+ \sqrt{4\alpha_2\frac{\alpha_1 - \alpha_2 - 1 + \sqrt{4\alpha_1 + (\alpha_1 - \alpha_2 - 1)^2}}{2 + \alpha_1 - \alpha_2 - 1 + \sqrt{4\alpha_1 + (\alpha_1 - \alpha_2 - 1)^2}} + \left(\alpha_2\frac{\alpha_1 - \alpha_2 - 1 + \sqrt{4\alpha_1 + (\alpha_1 - \alpha_2 - 1)^2}}{2 + \alpha_1 - \alpha_2 - 1 + \sqrt{4\alpha_1 + (\alpha_1 - \alpha_2 - 1)^2}} - \alpha_3 - 1\right)^2}\Bigg\}. \end{aligned} \quad [6]$$

### Analysis and (in)sensitivity

In this part of the supplementary material we showcase the generality of our results, based on the generalized model (Eq. (3)). We first show that increasing the concentration of one catalyst acting on a single step always results in a diminishing return of investment, so concentrating one catalyst in part of the hyphae leads to a lower overall harvest rate (Corollary 1). Then we show that increasing all catalyst concentrations along the multi-step process always yields economy of scale, so concentrating all catalysts in part of the hyphae results in a higher overall harvest rate (Corollary 2). This is complemented by looking at the limit behavior of the harvest rate at low and high catalyst concentrations for any number of catalysts. These results are all insensitive to parameter values. The effect of concentrating several (but not all) catalysts on the return of investment does depend on parameter values (Corollary 5).

**Concentrating a single catalyst: diminishing return of investment.** Recall that we use as notation for the conversion rates $r_1 = k_1\alpha_1$ and $r_i = \frac{k_i\alpha_ip_i}{K_i+p_i}$ $(i > 1)$. In equilibrium, we also use $\bar{r}_j = \frac{k_j\alpha_j\bar{p}_j}{K_j+\bar{p}_j}$ (and $\bar{r}_1 = r_1$) and have:

$$\bar{r}_i = l_{i+1}\bar{p}_{i+1} + \bar{r}_{i+1} \qquad 1 \leq i \leq n. \quad [7]$$

**Lemma 1.** *If the equilibrium conversion rate $\bar{r}_i$ from $P_i$ to $P_{i+1}$ $(i < n)$ changes by a factor $s$ and everything else stays the same, then the subsequent conversion rate $\bar{r}_{i+1}$ changes by a factor between 1 and $s$.*

*Proof.* This follows from the fact that the loss term $l_{i+1}p_{i+1}$ is linear in $p_{i+1}$ whilst the conversion term $r_{i+1} = \frac{k_{i+1}\alpha_{1+1}p_{i+1}}{K_{i+1}+p_{i+1}}$ is also increasing but has a horizontal asymptote. For $s > 1$ $(s < 1)$ this implies that $\bar{p}_{i+1}$ increases (decreases) and results in a larger (smaller) proportion of loss compared to conversion, thus dampening the impact of the change of the conversion rate $\bar{r}_i$ to the next $\bar{r}_{i+1}$. □

**Corollary 1.** *Increasing the concentration of one catalyst acting on a single step in the multi-step process leads to a diminishing return of investment. In particular, concentrating one catalyst in part of the hyphae (keeping the total amount of catalyst constant) leads to a lower overall harvest rate.*

*Proof.* Suppose that all catalyst concentrations are constant, except for one catalyst $A_j$ with concentration $q\alpha_j$ depending on $q$.

If $j = 1$, then the conversion rate $\bar{r}_1 = k_1q\alpha_1$ from $P_1$ to $P_2$ changes by a factor $q$. By lemma 1, each subsequent conversion rate will also change, with a factor that remains between 1 and $q$.

For $j > 1$, the conversion $\bar{r}_{j-1}$ from $P_{j-1}$ to $P_j$ will remain unchanged, the dependence of $A_j$ on $q$ only influences the process from $P_j$ onwards. The factor by which the conversion rate $\bar{r}_j = \frac{k_jq\alpha_j\bar{p}_j}{K_j+\bar{p}_j}$ changes lies between 1 and $q$, because although the catalyst concentration changes by a factor $q$, the equilibrium concentration $\bar{p}_j$ will change in the opposite direction. As before, by Lemma 1, the factor of change will remain between 1 and $q$ for subsequent conversion rates.

As the harvest rate is the last conversion rate, the change of the harvest rate is smaller than the change of the concentration of the catalyst $A_j$. □

So, by increasing the concentration of one catalyst $A_j$ by a factor $q$, the harvest rate will increase by a factor less than $q$. Similarly, for $q < 1$, the harvest rate will decrease less than the decrease of the catalyst concentration.

**Concentrating all catalysts: economy of scale.** Instead of changing one catalyst concentration, we will now change all catalyst concentrations by the same factor $q$. The enzyme concentrations are thus given by $q\boldsymbol{\alpha}$, with $q = 1$ the point of reference. We introduce notation $\bar{p}_i(q)$ for the corresponding equilibrium concentration of intermediate product $P_i$ and start with two lemmas.

**Lemma 2.** *Let $a, x > 0$, then $\frac{a+bx}{a+x}$ lies between 1 and $b$.*

*Proof.* We have $\frac{a+bx}{a+x} = b + \frac{1-b}{a+x}$ which implies that the fraction is below (for $b > 1$) or above (for $b < 1$) $b$. We also have $\frac{a+bx}{a+x} = 1 + \frac{(r-1)x}{a+x}$ which implies that the fraction is above (for $b > 1$) or below (for $b < 1$) 1. In summary, the fraction is between 1 and $b$. □

**Lemma 3.** *The equilibrium concentration $\bar{p}_i(q)$ with catalyst concentrations given by $q\boldsymbol{\alpha}$ lies between $\bar{p}_i(1)$ and $q^{i-1}\bar{p}_i(1)$.*

*Proof.* Proof by induction. For the initial step we need to show that $\bar{p}_2(q)$ lies between $\bar{p}_2(1)$ and $q\bar{p}_2(1)$. For $q = 1$, in equilibrium we have that the source term of $P_2$ equals the sum of the sink terms (Eq. (7)). Multiplying both sides by $q$ we obtain:

$$q(k_1\alpha_1) = q\left(l_2\bar{p}_2(1) + \frac{k_2\alpha_2\bar{p}_2(1)}{K_2 + \bar{p}_2(1)}\right). \quad [8]$$

Equilibrium for general $q$ implies:

$$k_1 q\alpha_1 = l_2\bar{p}_2(q) + \frac{k_2 q\alpha_2\bar{p}_2(q)}{K_2 + \bar{p}_2(q)}. \quad [9]$$

Combining these equations we get:

$$l_2(q\bar{p}_2(1) - \bar{p}_2(q))(K_2 + \bar{p}_2(1))(K_2 + \bar{p}_2(q)) = k_2 q\alpha_2 K_2(\bar{p}_2(q) - \bar{p}_2(1)). \quad [10]$$

So $\operatorname{sgn}(q\bar{p}_2(1) - \bar{p}_2(q)) = \operatorname{sgn}(\bar{p}_2(q) - \bar{p}_2(1))$. From this it follows that $\bar{p}_2(q)$ lies between $\bar{p}_2(1)$ and $q\bar{p}_2(1)$. In particular, $\bar{p}_2(1) < \bar{p}_2(q) < q\bar{p}_2(1)$ if $q > 1$ and $q\bar{p}_2(1) < \bar{p}_2(q) < \bar{p}_2(1)$ if $0 < q < 1$.

For the induction step, suppose that $\bar{p}_{m-1}(q)$ lies between $\bar{p}_{m-1}(1)$ and $q^{m-2}\bar{p}_{m-1}(1)$. In equilibrium for $q = 1$ we have:

$$q\frac{k_{m-1}\alpha_{m-1}\bar{p}_{m-1}(1)}{K_{m-1} + \bar{p}_{m-1}(1)} = q\left(l_m\bar{p}_m(1) + \frac{k_m\alpha_m\bar{p}_m(1)}{K_m + \bar{p}_m(1)}\right). \quad [11]$$

And in equilibrium for general $q$ we have:

$$\frac{k_{m-1}q\alpha_{m-1}\bar{p}_{m-1}(q)}{K_{m-1} + \bar{p}_{m-1}(q)} = l_m\bar{p}_m(q) + \frac{k_m q\alpha_m\bar{p}_m(q)}{K_m + \bar{p}_m(q)}. \quad [12]$$

Combining these equations we get:

$$l_m(q\bar{p}_m(1)(K_{m-1} + \bar{p}_{m-1}(q)) - \bar{p}_m(q)(K_{m-1} + \bar{p}_{m-1}(1))(K_m + \bar{p}_m(1))(K_m + \bar{p}_m(q)) = k_m q\alpha_m K_m(\bar{p}_m(q) - \bar{p}_m(1)). \quad [13]$$

So $\operatorname{sgn}(q\bar{p}_m(1)(K_{m-1}+\bar{p}_{m-1}(q)) - \bar{p}_m(q)(K_{m-1}+\bar{p}_{m-1}(1))) = \operatorname{sgn}(\bar{p}_m(q) - \bar{p}_m(1))$. From this it follows that $\bar{p}_m(q)$ lies between $\bar{p}_m(1)$ and $q\bar{p}_m(1)\frac{K_{m-1}+\bar{p}_{m-1}(q)}{K_{m-1}+\bar{p}_{m-1}(1)}$. By assumption, $\bar{p}_{m-1}(q)$ lies between $\bar{p}_{m-1}(1)$ and $q^{m-2}\bar{p}_{m-1}(1)$, which after application of Lemma 2 implies that $\frac{K_{m-1}+\bar{p}_{m-1}(q)}{K_{m-1}+\bar{p}_{m-1}(1)}$ lies between 1 and $q^{m-2}$. Thus we find that $\bar{p}_m(q)$ lies between $\bar{p}_m(1)$ and $q^{m-1}\bar{p}_m(1)$ for all $m$. In particular, $\bar{p}_m(1) < \bar{p}_m(q) < q^{m-1}\bar{p}_m(1)$ if $q > 1$ and $q^{m-1}\bar{p}_m(1) < \bar{p}_m(q) < \bar{p}_m(1)$ if $0 < q < 1$. □

**Corollary 2.** *The harvest rate $h(q\boldsymbol{\alpha})$ where all catalyst concentrations are multiplied by the same factor $q$ lies between $qh(\boldsymbol{\alpha})$ and $q^n h(\boldsymbol{\alpha})$, where $n$ is the number of steps in the process. So increasing the concentration of all catalysts together leads to an increased return of investment: economy of scale. In particular, concentrating all catalysts in part of the hyphae (keeping the total amount of catalyst constant) results in a higher overall harvest rate.*

*Proof.* The harvest rate (Eq. (5)) with variable catalyst concentrations is given by $h(q\boldsymbol{\alpha}) = \frac{k_n q\alpha_n\bar{p}_n(q)}{K_n+\bar{p}_n(q)}$, which is a monotonically increasing function of $\bar{p}_n(q)$. Substituting $\bar{p}_n(q) = \bar{p}_n(1)$ yields that $h(q\boldsymbol{\alpha})$ is bound by $\frac{k_n q\alpha_n\bar{p}_n(1)}{K_n+\bar{p}_n(1)} = qh(\boldsymbol{\alpha})$. Substituting $\bar{p}_n(q) = q^{n-1}\bar{p}_n(1)$ yields that $h(q\boldsymbol{\alpha})$ is bound by $\frac{k_n q\alpha_n q^{n-1}\bar{p}_n(1)}{K_n+q^{n-1}\bar{p}_n(1)} = q^n\frac{k_n\alpha_n\bar{p}_n(1)}{K_n+q^{n-1}\bar{p}_n(1)}$ which is a stronger bound than $q^n h(\boldsymbol{\alpha})$. □

**Asymptotics for low and high catalyst concentration.** We assume that the concentration of each catalyst $A_i$ is either $\alpha_i$ (a constant) or $q\alpha_i$ (dependent on the catalyst concentration parameter $q$). Before in this SI, we restricted our attention to varying the concentration of either a single or all catalysts, here we vary any number of catalysts.

Recall that the harvest rate equals the last conversion rate of the multi-step process in equilibrium (Eq. (5)). For low $q$, we show that the harvest rate can be approximated by a power function with exponent $m$ equal to the number of catalyst concentrations that are $q$-dependent. For this we need a preparatory Lemma.

**Lemma 4.** *Suppose $1 < q \ll 1$. If we are only interested in the harvest rate, then we can make the following reductions within the multi-step process:*

1. *If $r_{j-1} = c_{j-1}$ does not depend on $q$ and $r_j = \frac{k_j q\alpha_j p_j}{K_j+p_j}$ does, we can replace the two steps $P_{j-1} \to P_j \to P_{j+1}$ by $P_{j-1} \to P_{j+1}$ with conversion rate $\frac{k_j\alpha_j c_{j-1}q}{l_jK_j+c_{j-1}}$.*
2. *If $r_{j-1} = c_{j-1}q^d$ (with $d \geq 1$) and $r_j = \frac{k_j\alpha_j p_j}{K_j+p_j}$ does not depend on $q$, we can replace the two steps $P_{j-1} \to P_j \to P_{j+1}$ by $P_{j-1} \to P_{j+1}$ with conversion rate $\frac{k_j\alpha_j c_{j-1}q^d}{l_jK_j+k_j\alpha_j}$.*
3. *If $r_{j-1} = c_{j-1}q^d$ (with $d \geq 1$) and $r_j = \frac{k_j q\alpha_j p_j}{K_j+p_j}$ also depends on $q$, we can replace the two steps $P_{j-1} \to P_j \to P_{j+1}$ by $P_{j-1} \to P_{j+1}$ with conversion rate $\frac{k_j\alpha_j c_{j-1}q^{d+1}}{l_j}$.*

*Proof.* Each of the reductions follows from application of Eq. (7). Specifically:

1. For $0 < q \ll 1$, we obtain $\bar{p}_j \approx \frac{c_{j-1}}{l_j}$ so that $r_j \approx \frac{k_j q\alpha_j \frac{c_{j-1}}{l_j}}{K_j+\frac{c_{j-1}}{l_j}} = \frac{k_j\alpha_j c_{j-1}q}{l_jK_j+c_{j-1}}$.
2. It holds that $\bar{p}_j < \frac{c_{j-1}q^d}{l_j} \ll 1$ so that $r_j \approx \frac{k_j\alpha_j\bar{p}_j}{K_j}$, leading to $p_j \approx \frac{c_{j-1}q^d}{l_j+\frac{k_j\alpha_j}{K_j}} = \frac{c_{j-1}K_jq^d}{l_jK_j+k_j\alpha_j}$ and finally $r_j \approx \frac{k_j\alpha_j c_{j-1}q^d}{l_jK_j+k_j\alpha_j}$.
3. It holds that $\bar{p}_j \lesssim \frac{c_{j-1}q^d}{l_j} \ll 1$ so $r_j \approx \frac{k_j q\alpha_j\bar{p}_j}{K_j} \approx \frac{k_j\alpha_j c_{j-1}q^{d+1}}{l_j}$.

□

The reductions of Lemma 4 are graphically depicted in Fig. S2 and are used to prove the following Corollary.

**Corollary 3.** *For $0 < q \ll 1$, it holds that $h \approx cq^m$, with $m$ the number of catalyst concentrations that depend on $q$, and $c$ a constant.*

*Proof.* For $m = 0$, none of the catalyst concentrations varies (with $q$), so the harvest rate does not depend on $q$ (and can be viewed as a power function with exponent 0). If $m \geq 1$ catalysts vary with $q$, let $A_j$ be the first such catalyst in the multi-step process, so $r_j$ depends on $q$. By repeated use of reduction 1. from Lemma 4, all conversion rates $r_1$ up to $r_j$ can be replaced by a single step that depends linearly on $q$. Subsequently, applying either reduction 2. or 3. of Lemma 4, each time the next conversion rate depends on $q$, another factor $q$ is picked up. So, regarding the equilibrium harvest rate, the entire multi-step process is equivalent to a single conversion rate that is a power function of $q$ with power $m$. □

Having treated the case where $q$ is very small, $0 < q \ll 1$, we now move to the opposite limiting case. For very high $q$, we will show that the harvest rate has a horizontal asymptote if not all catalysts vary with $q$ and has an oblique asymptote if all catalysts vary with $q$. For this, again, we first prove a Lemma.

**Lemma 5.** *Suppose $q \gg 1$. If we are only interested in the harvest rate, then we can make the following reductions within the multi-step process:*

1. *If $r_1 = k_1q\alpha_1$ depends on $q$ and $r_2 = \frac{k_2\alpha_2p_2}{K_2+p_2}$ does not, we can replace the two steps $P_1 \to P_2 \to P_3$ by $P_1 \to P_3$ with conversion rate $k_2\alpha_2$.*
2. *If $r_1 = k_1q\alpha_1$ depends on $q$ and so does $r_2 = \frac{k_2q\alpha_2p_2}{K_2+p_2}$, we can replace the two steps $P_1 \to P_2 \to P_3$ by $P_1 \to P_3$ with conversion rate $q\min(k_1\alpha_1, k_2\alpha_2)$.*
3. *If $r_{j-1}$ does not depend on $q$ and $r_j = \frac{k_jq\alpha_jp_j}{K_j+p_j}$ does, we can replace the two steps $P_{j-1} \to P_j \to P_{j+1}$ by $P_{j-1} \to P_{j+1}$ with conversion rate $r_{j-1}$.*

*Proof.* Again we make use of Eq. (7) in each of the cases:

1. It holds that $r_2 < k_2\alpha_2$ so that $\bar{p}_2 > \frac{k_1q\alpha_1}{k_2\alpha_2+l_2} \to \infty$ as $q \to \infty$. In this case, $r_2 \to k_2\alpha_2$.

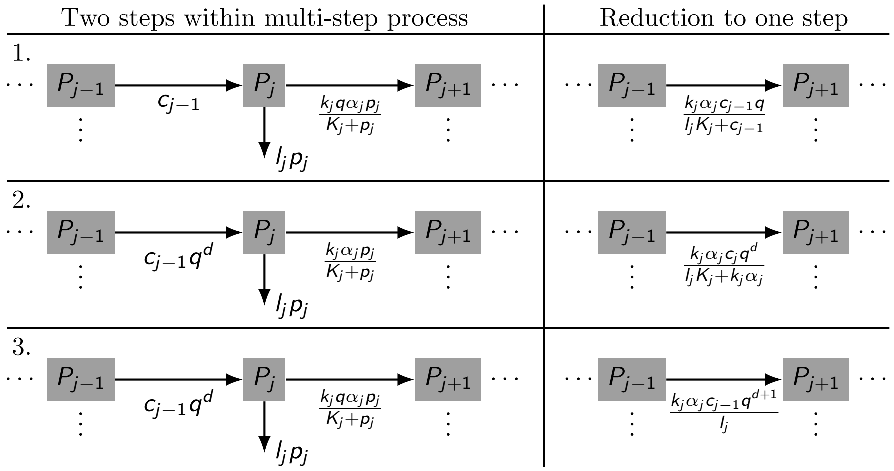


**Fig. S2.** Graphical representation of the reductions 1., 2. and 3. of the multi-step process for $0 < q \ll 1$, as treated in Lemma 4.

2. We have $k_1\alpha_1 = \frac{k_2\alpha_2\bar{p}_2}{K_2+\bar{p}_2} + \frac{l_2\bar{p}_2}{q}$. Suppose $k_1\alpha_1 < k_2\alpha_2$, then $\bar{p}_2 < \frac{k_1\alpha_1 K_2}{k_2\alpha_2 - k_1\alpha_1}$ which implies that $\frac{l_2\bar{p}_2}{q} \to 0$ as $q \to \infty$. So $\bar{p}_2 \to \frac{k_1\alpha_1 K_2}{k_2\alpha_2-k_1\alpha_1}$ and $r_2 \to \frac{k_2 q\alpha_2 k_1\alpha_1 K_2}{K_2(k_2\alpha_2-k_1\alpha_1)+k_1\alpha_1 K_2} = qk_1\alpha_1$. Else suppose $k_1\alpha_1 > k_2\alpha_2$, from $k_1\alpha_1 < k_2\alpha_2 + \frac{l_2\bar{p}_2}{q}$ we obtain $\bar{p}_2 > \frac{q(k_1\alpha_1-k_2\alpha_2)}{l_2} \to \infty$ as $q \to \infty$ so that $r_2 \to \frac{k_2 q\alpha_2\bar{p}_2}{\bar{p}_2} = qk_2\alpha_2$. Taken together, this yields the conversion rate $q\min(k_1\alpha_1, k_2\alpha_2)$ for the reduction to one step.

3. It holds that $r_{j-1} > r_j = \frac{k_j q\alpha_j\bar{p}_j}{K_j+\bar{p}_j}$ so $\bar{p}_j < \frac{r_{j-1}K_j}{k_j q\alpha_j - r_{j-1}} \ll 1$ for $q \gg 1$. So $r_j \approx \frac{k_j q\alpha_j\bar{p}_j}{K_j}$ and from this $\bar{p}_j \approx \frac{r_{j-1}}{l_j+\frac{k_j q\alpha_j}{K_j}} = \frac{r_{j-1}K_j}{l_j K_j + k_j q\alpha_j} \approx \frac{r_{j-1}K_j}{k_j q\alpha_j}$ for $q \gg 1$ from which $r_j \approx r_{j-1}$.

□

The reductions of Lemma 5 are graphically depicted in Fig. S3 and are used to prove the following Corollary.

**Corollary 4.** *If not all catalyst concentrations vary with $q$, then the harvest rate $h$ has a horizontal asymptote for $q \to \infty$. If all catalyst concentrations vary with $q$, $h$ has an oblique asymptote with slope $\min(k_1\alpha_1, k_2\alpha_2, \ldots, k_n\alpha_n)$.*

*Proof.* If the first catalyst concentration varies with $q$, we can either use reduction 1. or 2. from Lemma 5. In case of applying reduction 2., we return to the original situation where the first catalyst concentration varies with $q$ again. There are now two possibilities. If all catalyst concentrations vary with $q$, repeated use of reduction 2. yields that the harvest rate of the multi-step process for $q \to \infty$ is equivalent to a single reaction with conversion rate $q\min(k_1\alpha_1, k_2\alpha_2, \ldots, k_n\alpha_n)$: an oblique asymptote. If not all catalysts vary with $q$, eventually reduction 1. can be applied. Now the first catalyst concentration does not depend on $q$. In this case, application of reduction 3. to any conversion rate with $q$-dependence results in a reduced multi-step process without $q$-dependence. Thus, we find that if not all catalyst concentrations vary with $q$, the equilibrium harvest rate has a horizontal asymptote. □

**Corollary 5.** *If several (not all) catalyst concentrations vary with $q$, depending on parameter values this can lead to either increasing return (economy of scale) or a diminishing return of investment.*

*Proof.* For $0 < q \ll 1$ there is an increasing return by Corollary 3. For $q \gg 1$ there is a diminishing return by Corollary 4. □

Our asymptotic analysis was graphically summarized in Fig. 3 (main article). By corollary 5, it depends on the parameter values whether concentrating two catalysts in the three-step amylolytic model is beneficial or not. To illustrate this, we vary the parameter $k_1$ while keeping all other parameters except $q$ constant (and still equal to one). Next to the value $k_1 = 1$ we analyzed in the article, we here analyze $k_1 = 0.1$ and $k_1 = 10$.

For $k_1 = 0.1$, the first conversion rate $r_1$ is ten times smaller than that of Fig. 1. Thus, saturation effects in subsequent conversion rates play a smaller role. Because of this, the range of increasing return of investment in the green scenario - with two varying catalyst concentrations - dominates, see Fig. S4B.

On the contrary, for $k_1 = 10$ the first conversion rate $r_1$ is ten times larger than that of Fig. 1. Thus, saturation effects in subsequent conversion rates play a larger role. Because of this, the range of diminishing return of investment in the green scenario dominates (Fig. S5B).

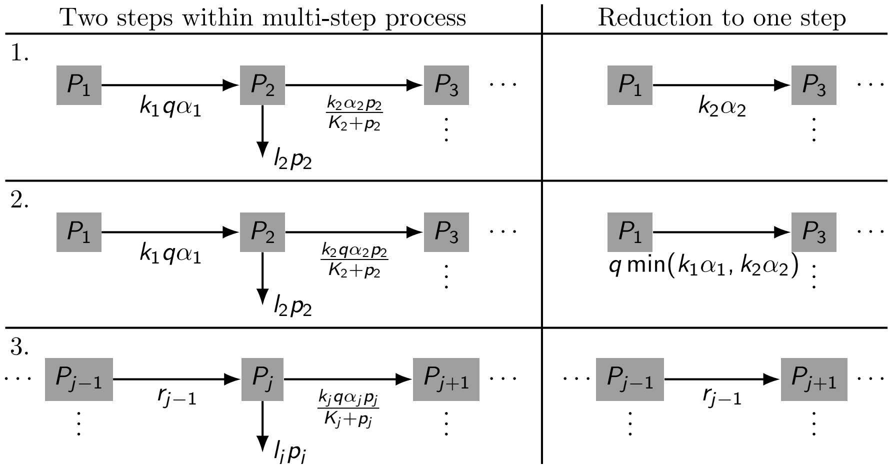


**Fig. S3.** Graphical representation of the reductions 1., 2. and 3. of the multi-step process for $q \gg 1$, as treated in Lemma 5.

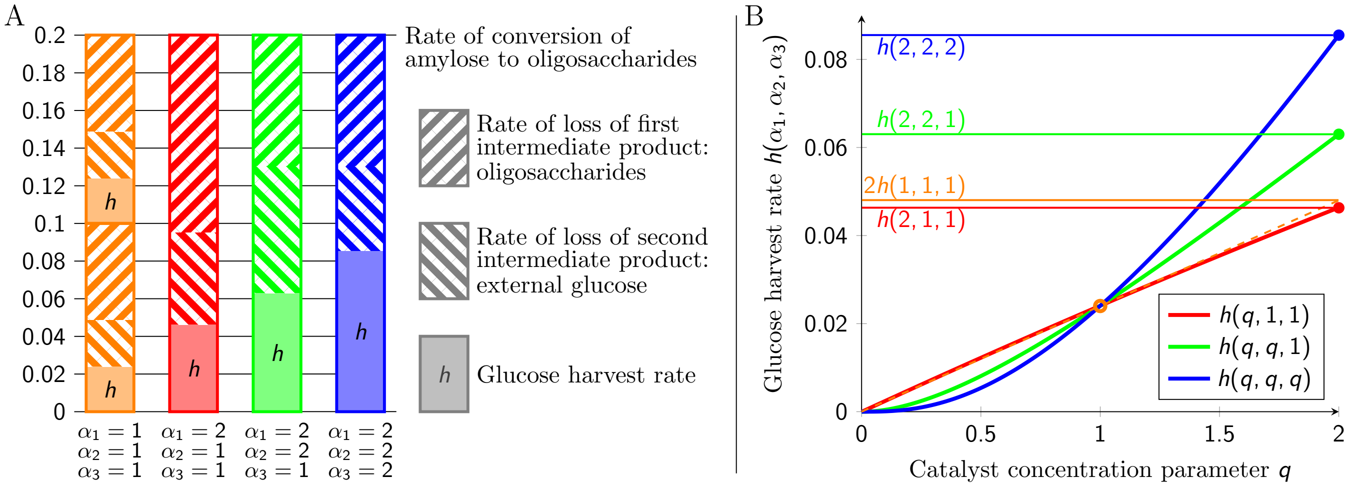


**Fig. S4.** (A and B) Glucose harvest rate for $k_1 = 0.1$ instead of the default value $k_1 = 1$ (confer Fig. 1). As before, $k_2 = k_3 = K_2 = K_3 = l_2 = l_3 = 1$. Relatively low $k_1 = 0.1$ results in a low flux through the multi-step process and low equilibrium concentrations $\bar{p}_2$ and $\bar{p}_3$, so that saturation in the conversion rates $r_2$ and $r_3$ hardly plays a role. This is beneficial for the harvest rates of concentrating catalysts compared to the benchmark: even concentrating one catalyst (red scenario) approximates the fifty-fifty benchmark (orange). Concentrating two catalysts (green scenario) results in a higher average harvest rate than fifty-fifty: $0.032 > h(1,1,1) \approx 0.024$. (B) Here, the increasing slope part of the sigmoidal green curve dominates, corresponding to increasing return of investment.

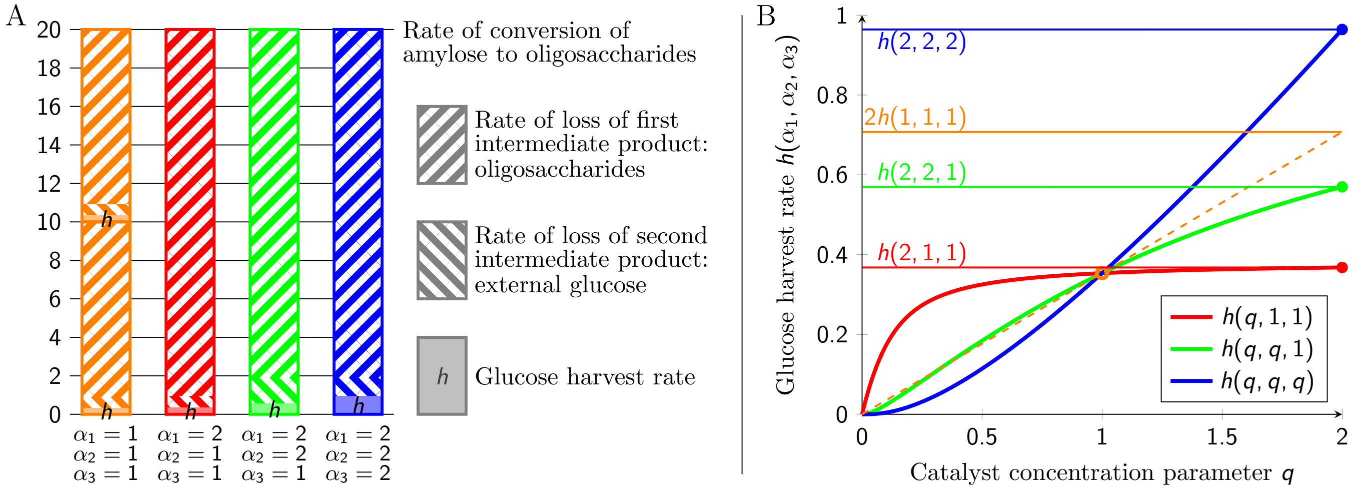


**Fig. S5.** (A and B) Glucose harvest rate for $k_1 = 10$ instead of the default value $k_1 = 1$ (confer Fig. 1). As before, $k_2 = k_3 = K_2 = K_3 = l_2 = l_3 = 1$. Relatively high $k_1 = 10$ results in high equilibrium concentrations $\bar{p}_2$ and $\bar{p}_3$. If only part of the catalysts is concentrated this leads to congestion and relative high loss of intermediate products due to saturation in the conversion rates. This is detrimental for concentrating catalysts: even concentrating all catalysts (blue scenario) results in a relatively minor advantage over fifty-fifty. Here, concentrating two catalysts (green scenario) results in a lower average harvest rate than fifty-fifty: $0.28 < h(1, 1, 1) \approx 0.35$. (B) Now the decreasing slope part of the sigmoidal green curve dominates, corresponding to diminishing return of investment.

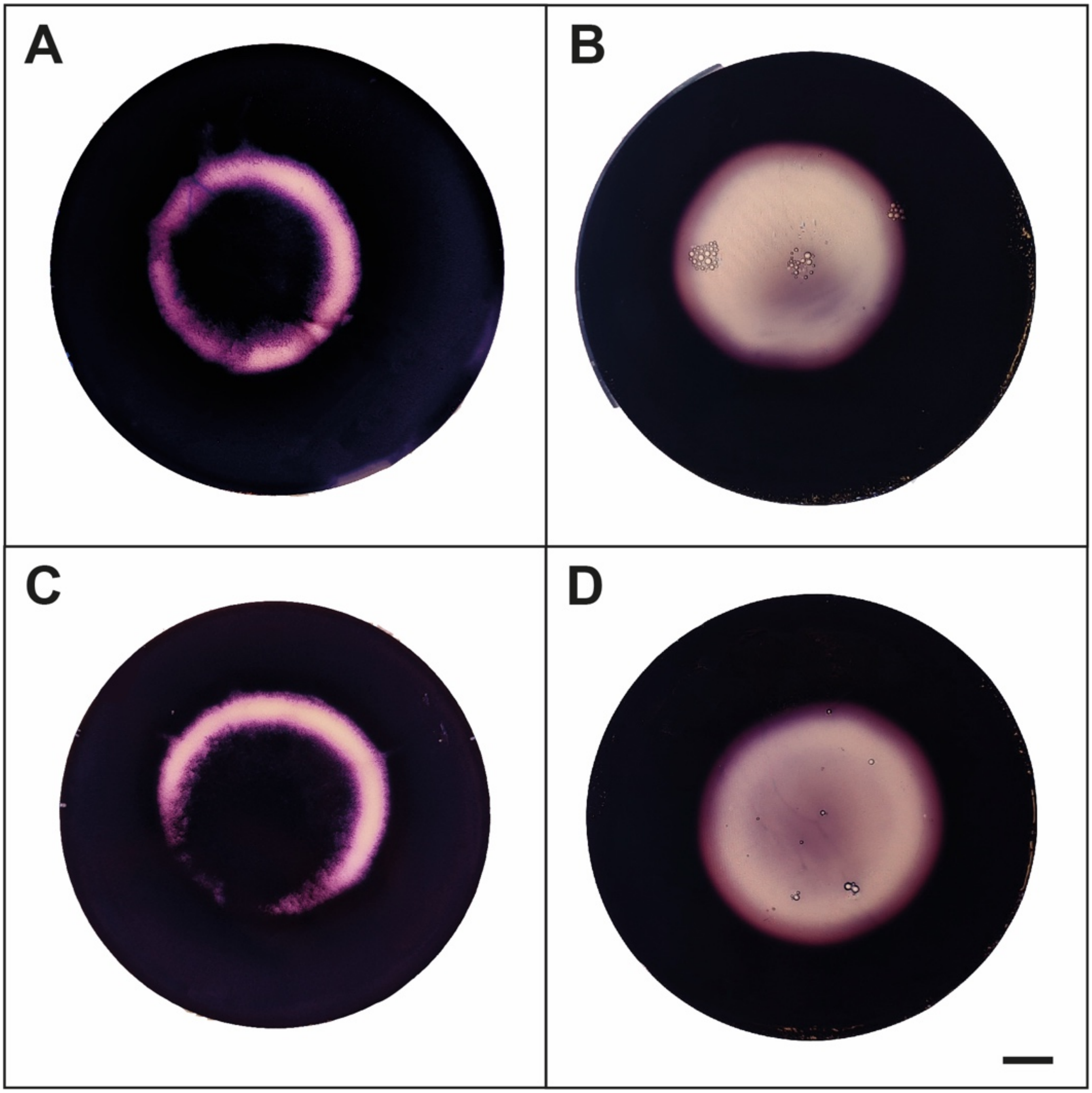


**Fig. S6.** MM agar medium supplemented with $2\%$ starch stained with Lugol after incubation with $5$-day old *A. niger* N402 (A, B) or N402$\Delta$*hexA* sandwich colonies (C, D), for $8$ h (A, C) and $24$ h (B, D). Scale bar represents 1 cm. Lugol can only bind the helix structure of starch, but not its degradation products, that lack this structure. The difference in harvest rate observed in the experiment is thus not due to a difference in conversion rate from amylose to oligosaccharides, but must be due to efficiency of heterogeneity in subsequent conversion steps.
